# Dose–Response Alignment Does Not Inherently Enhance Information Transmission in Signaling Pathways

**DOI:** 10.64898/2026.06.10.731507

**Authors:** David Gordillo-Alaniz, Eugenio Azpeitia

## Abstract

Understanding how signaling pathways reliably transmit information is fundamental to explaining cellular decision-making. Several studies have suggested that Dose–Response Alignment (DoRA) —the overlapping of dose–response curves between receptor occupancy and downstream responses— enhances information transmission by ensuring a linear mapping between occupancy and response. Here, we challenge this intuition using analytical and numerical analyses of a basic signaling mechanism. Our results show that DoRA, linearity and information transmission are fundamentally dissociable properties. We find that occupancy–response linearity is normalization-independent, whereas the measured degree of alignment depends on the normalization scheme used to compare dose–response curves. Specifically, normalization by maximal attained responses preferentially reflects occupancy–response linearity, whereas normalization by total response capacity preserves response dynamic range. Accordingly, under the first normalization, perfect alignment reflects linear occupancy–response mappings, whereas under the second normalization alignment can remain imperfect because it does not exclusively reflect linearity. Importantly, neither perfect alignment nor linearity necessarily maximize information transmission; instead, information transmission can increase through larger dynamic range or reduced noise. Our results show that efficient information transmission cannot be inferred from dose–response alignment or linearity alone, but emerges from the balance between dynamic range, biochemical noise, and input–output mapping.

## Introduction

Cells must continually sense and interpret cues from their surroundings to survive, grow, and carry out their physiological functions [1]. Reliable acquisition of environmental information is therefore fundamental to living systems [2]. At the cellular level, this task is carried out primarily by signaling pathways, which detect extracellular stimuli and propagate this information through cascades of biochemical interactions [3]. The more accurately a cell can detect environmental changes, the more effectively it can adjust its internal state and produce an appropriate response, ultimately enhancing its survival and fitness [4, 5].

Signaling pathways operate under multiple constraints that limit the precision with which cells extract information from their environment [2, 6]. These constraints include the stochastic nature of the underlying biochemical reactions and cell-to-cell variability in the abundance of signaling components [7, 8, 9, 10]. Such noise and variability are inherent to all signaling pathways and can degrade the fidelity with which external cues are detected and transmitted [11, 12, 13]. Because these constraints are unavoidable, considerable effort has been devoted to understanding how signaling pathways are organized to overcome, or at least mitigate, their effects.

Several molecular and dynamical mechanisms have been proposed as potential contributors to improved information transmission [14, 15]. Among these mechanisms, particular attention has been given to those that produce linear relationships between receptor occupancy and downstream responses [16, 17, 18]. A linear *occupancy–response* relationship implies that changes in the detected stimulus are translated into proportional changes in the cellular downstream response. Such proportionality has been proposed to enhance information transmission in signaling pathways through different mechanisms, including: (a) allowing response magnitude to reflect stimulus intensity across a broad input range, (b) reducing distortions in signal propagation arising from nonlinear reactions within signaling cascades, and (c) helping to delay early signal saturation [19, 20]. Importantly, proportionality has been identified in key regulatory networks, such as the NF-*κ*B and MAPK pathways, where feedback loops and signaling dynamics appear to maintain a broad dynamic range and high-resolution output [21, 22, 23, 16]. Together, these observations suggest that linear occupancy–response relationships may provide functional regimes for preserving stimulus information across signaling pathways.

Motivated by the need to experimentally characterize these proportional input–ouput relationships, several quantitative approaches have been developed to assess the degree of linearity in signaling pathways [24, 25]. In particular, it has been proposed that linear *occupancy–response* relationships can be inferred by measuring the extent to which the dose–response curves of receptor occupancy and downstream responses overlap, a property known as *dose–response alignment* (DoRA) [19, 26, 24]. In fact, empirical studies comparing receptor occupancy and downstream responses have reported that diverse signaling pathways, including the pheromone response pathway in *Saccharomyces cerevisiae* [19, 26, *24] and conserved mammalian pathways such as Wnt, ERK, and TGF–β*, operate in a DoRA regime [18]. However, whether *dose–response alignment* reflects underlying linearity of the pathway and contributes to accurate information transmission has not been formally established.

Importantly, the relationship between DoRA, *occupancy–response* linearity, and information transmission is not straightforward and is affected by several factors. First, dose–response curves of receptor occupancy and downstream responses typically differ in magnitude. For this reason, their comparison requires normalization to a common scale [27]. Different normalization schemes have been used in the literature, including scaling relative to the maximal observed response or to the total available response. This methodological choice may therefore influence whether DoRA reflects an underlying linear occupancy–response relationship. However, how normalization choices affect the inferred presence and apparent strength of DoRA, and its ability to reflect linear occupancy–response relationships, has not been explicitly addressed.

Second, information transmission depends not only on occupancy–response alignment, but also on other system properties such as noise and response dynamic range, which have been shown to exert opposing effects on signaling fidelity [28, 29]. Because these properties are jointly determined by underlying biochemical parameters, improvements in one aspect may come at the expense of others, reflecting inherent trade-offs in signaling pathway design. Nevertheless, how noise and response dynamic range modulate the relationship between DoRA, linearity, and information transmission remains poorly understood.

In this work, we use mathematical modeling and computational simulations to investigate the relationship between dose–response alignment (DoRA), occupancy–response linearity, and information transmission. We show that this relationship cannot be reduced to a simple equivalence, but instead depends on both the normalization procedures used to define and compare dose–response curves and on the noise and response dynamic range of the system. We specifically analyze how different normalization schemes affect the detection and quantification of DoRA and its relationship to underlying occupancy–response linearity. Then, we quantify how changes in noise and response dynamic range influence the relationship between DoRA, occupancy–response linearity, and information transmission. Together, these analyses allow us to assess whether increased alignment enhances information acquisition and how it interacts with other properties of signaling systems.

## The model

We used a minimal model of a cellular signaling pathway, in which an inactive receptor (*X*) binds to a stimulus molecule (*S*) and transitions to an occupied (active) state (*X*\*). The occupied receptor then promotes the activation of a downstream response protein (*Y→Y* *). (Fig. 1A) (see Methods and SI 1.1). We assumed fixed total concentrations of receptor (*X*_*T*_ = *X* + *X*\*) and response protein (*Y*_*T*_ = *Y* + *Y* *). Parameters *k*_1_ and *k*_2_ describe receptor–signal association and dissociation, *k*_3_ and *k*_5_ basal and receptor-mediated response activation, and *k*_4_ response deactivation. To isolate the effect of the receptor–response coupling, we set *k*_3_ = 0. Under this condition, the half-saturation constants *K*_*X*_ = *k*_2_*/k*_1_ and *K*_*Y*_ = *k*_4_*/k*_5_ determine the shape of dose–response curves (see SI 1.2).

**Figure 1:**
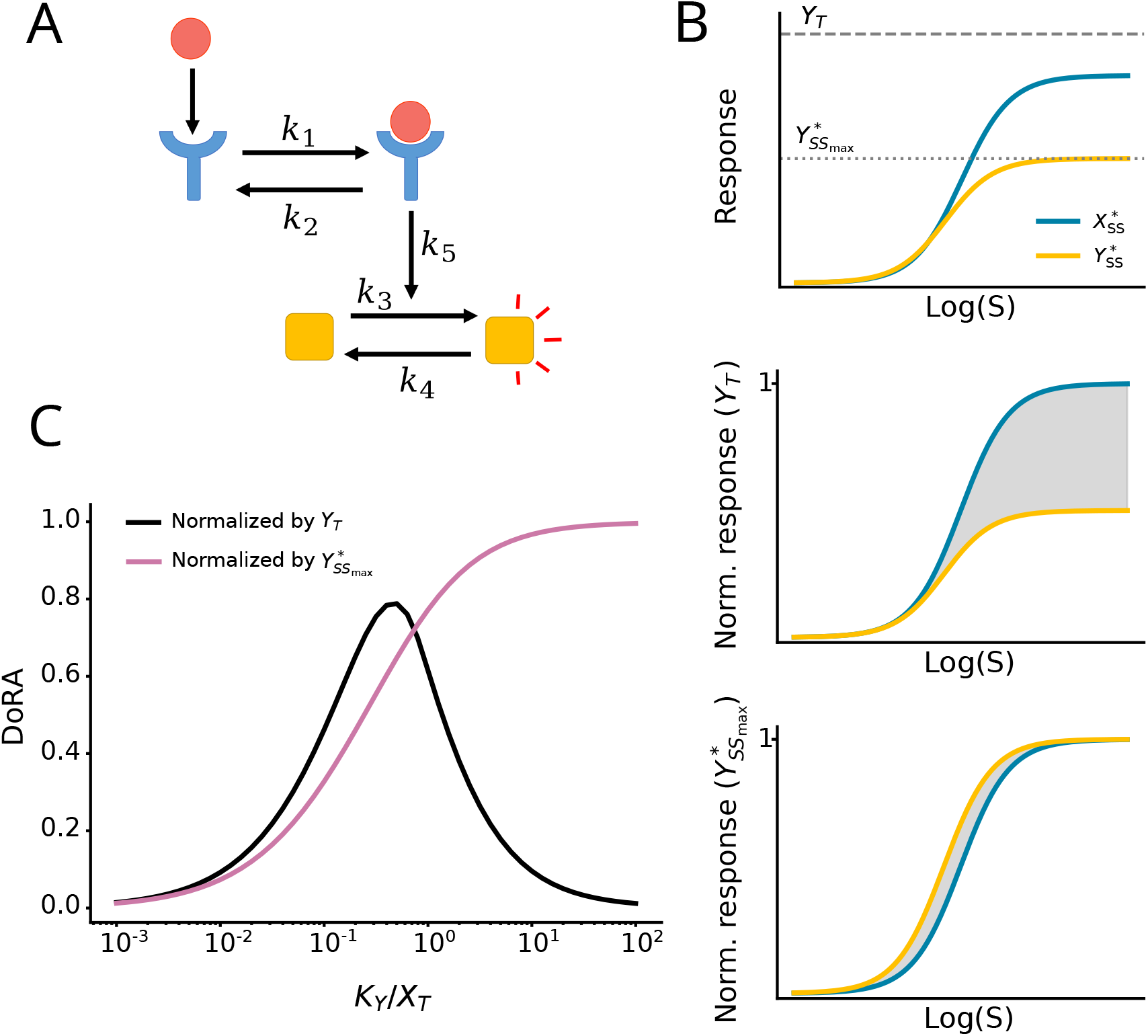
Model and dose–response alignment (DoRA). **A**, Schematic representation of our model of a signaling pathway. Parameters *k*_1_ and *k*_2_ represent receptor association and dissociation, respectively, whereas *k*_3_ and *k*_4_ represent activation of the downstream response, and *k*_5_ its deactivation. **B**, Effect of normalization on dose–response alignment (DoRA). **Top**, Unnormalized receptor occupancy and down-stream response dose–response curves. Horizontal lines indicate *Y*_*T*_ and 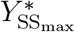 . **Middle**, Curves normalized by total available protein. **Bottom**, Curves normalized by the maximal observed response, for which both curves range from 0 to 1. Gray shading indicates the area between the receptor occupancy and downstream response curves, a measure of dose–response alignment. **C**, DoRA values obtained at different *K*_*Y*_ */X*_*T*_ ratios for both normalization schemes.

This signaling mechanism is particularly relevant because it is widely observed in common signaling pathways, including GTPase cycles in different organisms [30]. Moreover, previous studies have reported conflicting results regarding whether this mechanism produces DoRA or not [23, 24, 25].

## Results

### Dose–response alignment depends on the normalization scheme

In order to analyze how different normalization schemes affect dose–response alignment (DoRA), we constructed dose–response curves by computing the steady states of the occupied receptor 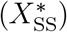 and downstream response 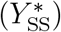 across 100 logarithmically spaced stimulus concentrations spanning three orders of magnitude around *K*_*X*_ = *k*_2_*/k*_1_, i.e *S* ∈ [10^*−*3^*K*_*X*_, 10^3^*K*_*X*_] (see Methods and SI 1.3). Within this signal range, receptor occupancy increased from nearly minimal 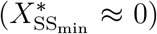 to almost complete saturation 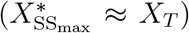. Before calculating DoRA, the dose–response curves were normalized employing two schemes commonly used in experimental analyses: (i) relative to the total available response protein (*X*_*T*_ or *Y*_*T*_) or (ii) relative to the maximal observed response 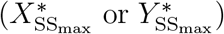 (Fig. 1B) (see Methods and SI 1.4). For the receptor, since 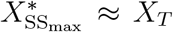, normalizing by *X*_*T*_ and 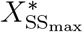 is effectively equivalent, therefore, the resulting normalized curve always represents a fractional response.

We then quantified DoRA from the normalized steady-state dose–response curves using a curve-distance metric based on previous formulations [25], which we rescaled such that a value of 1 corresponds to perfect alignment and 0 to maximal mismatch (see Methods and SI 1.5). In parallel, we analyzed the corresponding transfer function *y* = *f* (*x*) to determine conditions required for perfect alignment (see Methods and SI 1.5). Both metrics yielded qualitatively consistent results; we therefore present the main analyses using the curve-distance metric.

We found that the key parameter modulating the alignment between receptor occupancy and the downstream response is the ratio *K*_*Y*_ */X*_*T*_ (see Methods and SI 1.10 and 1.11). Importantly, the magnitude of DoRA depended strongly on the normalization scheme employed. When downstream responses were normalized by 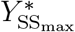, increasing *K*_*Y*_ */X*_*T*_ caused the downstream response curve to converge toward the receptor occupancy curve, with DoRA approaching 1 as *K*_*Y*_ */X*_*T*_*→* ∞ (Fig. 1C; pink line) (see SI 1.10). In contrast, normalization by *Y*_*T*_ prevented complete alignment. Under this scheme, maximal DoRA ≈ 0.8 occurred at ratio *K*_*Y*_ */X*_*T*_ ≈ 0.4622, when *K*_*Y*_ and *X*_*T*_ were of comparable magnitude (Fig. 1C; black line) (see SI 1.11). These results demonstrate that dose–response alignment depends on the normalization scheme used to normalize the downstream response, rather than arising solely from the intrinsic biochemical properties of the system.

### DoRA detection of linear occupancy–response relationships depends on the normalization scheme

Because *dose–response alignment* (DoRA) has been proposed to reflect the linearity of the system’s occupancy–response relationship [19]. Yet, because DoRA values depend on the normalization procedure, we next asked whether normalization also affects the relationship between DoRA and occupancy–response linearity. To address this question, we first characterized the occupancy–response relationship independently of normalization using the elasticity coefficient, defined as 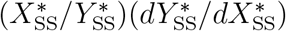, a scale-invariant measure of input–output proportionality that equals unity under locally linear dynamics [31, 32](see Methods and SI 1.6).

In the regime *K*_*Y*_ */X*_*T*_*→* ∞, the system’s occupancy–response relationship approaches linearity. Notably, when the response is normalized by 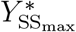, DoRA faithfully reflects this linear regime, approaching a value of 1. Under this normalization, plotting 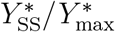 against normalized receptor occupancy produces curves that closely follow the identity line, as expected from a linear input–output relationship (Fig. 2A). In contrast, normalization by *Y*_*T*_ does not preserve this correspondence. In this case, even the *K*_*Y*_ */X*_*T*_ ratios associated with the highest DoRA values do not fall within the regime of near-linear behavior, as indicated by their deviation from the identity line (Fig. 2B). These results show that, when the response is normalized by 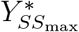, DoRA accurately reflects occupancy–response linearity. In contrast, when the response is normalized by *Y*_*T*_, high DoRA values do not necessarily correspond to a nearly linear occupancy–response relationship. Consequently, DoRA is not intrinsically equivalent to occupancy–response linearity. Rather, its apparent association with linearity emerges or disappears depending on the normalization used to represent the same underlying dynamics.

**Figure 2:**
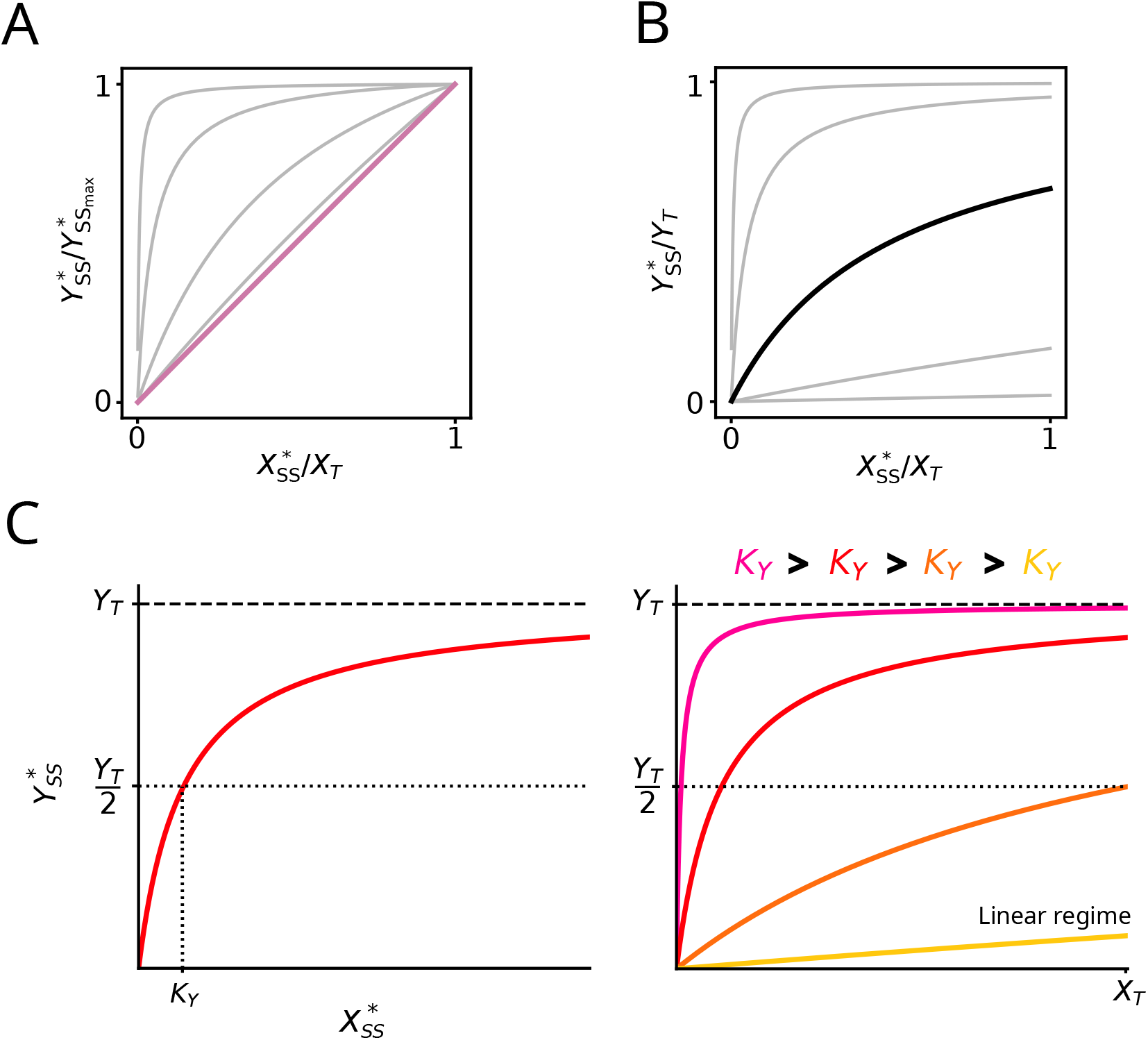
Relationship between DoRA, occupancy–response linearity, and normalization. Normalized occupancy–response curves, (**A**) 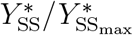 and (**B**) 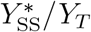, as a function of 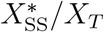. **A**, curves approach the identity line (pink line) as *K*_*Y*_ */X*_*T*_ increases, corresponding to increased occupancy–response linearity and higher DoRA under this normalization scheme. **B**, curves become progressively flatter as *K*_*Y*_ */X*_*T*_ increases. Under this normalization scheme, the curve yielding the maximal DoRA value (black curve) occurs outside the regime of near-linear occupancy–response behavior. **C**, Steady-state occupancy–response relationship underlying panels **A** and **B. Left**, Downstream response 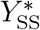 as a function of receptor occupancy 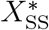. **Right**, Increasing *K*_*Y*_ (and therefore the *K*_*Y*_ */X*_*T*_ ratio) progressively linearizes the occupancy–response relationship while reducing the response range.

We next analyzed the steady-state expression of the downstream response as a function of receptor occupancy (see Methods and SI 1.2) to determine why the linear regime is only revealed under the 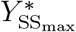 normalization scheme. The steady-state response can be written in the form of a Michaelis–Menten function of receptor occupancy (Fig. 2C; left),

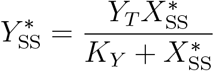

This expression directly defines the underlying occupancy–response relationship and shows that the downstream response becomes approximately linear when *K*_*Y*_ greatly exceeds the range of possible receptor occupancy values (Fig. 2C; right). Since 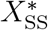 is bounded above by *X*_*T*_, the condition *K*_*Y*_ ≫ *X*_*T*_ ensures that *K*_*Y*_ ≫ 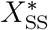 across the full activation range. In particular, in the regime *K*_*Y*_ */X*_*T*_ → ∞, the downstream response approaches the linear form,

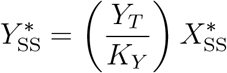

which corresponds to a linear occupancy–response relationship with proportionality factor *Y*_*T*_ */K*_*Y*_, but with a strongly reduced response magnitude and limited dynamic range.

This linear regime is revealed under normalization by 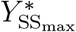 because the reduced response range characteristic of this regime is rescaled to span most of the normalized response range. As a result, DoRA values approach 1. In contrast, normalization by *Y*_*T*_ preserves the absolute response scale and does not rescale the dynamic range according to the observed response amplitude. Consequently, the approximately linear regime occupies only a small portion of the normalized response range, does not produce maximal curve alignment, and is therefore not reflected by DoRA.

Linear occupancy–response relationships have been associated with enhanced information transmission in signaling pathways [19, 24]. However, reduced response ranges can limit information transmission by compressing the range of distinguishable responses, while low response levels may increase the relative impact of molecular noise [28]. Here, we find that the normalization scheme yielding maximal DoRA values (i.e., normaliza-tion by 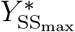) captures an approximately linear occupancy–response relationship, but at the cost of a reduced response range. In contrast, normalization by *Y*_*T*_ yields lower DoRA values, corresponding to a regime in which the near-linear behavior is not captured by the normalized representation, while preserving a broader dynamic range. These results raise the question of whether DoRA, under any normalization scheme, can serve as a reliable proxy for the information transmission capacity of the system, as previously suggested [19, 23, 24, 25].

### Normalization Shapes the Relationship Between DoRA and Information Transmission

To investigate how normalization influences the relationship between DoRA and information transmission, and whether occupancy–response linearity is associated with information transmission, we simulated the model using the Gillespie algorithm, which accounts for the intrinsic stochasticity of chemical reactions and generates statistically exact system trajectories [33, 34](see Methods and SI 1.7). Because stochastic steady states are characterized by probability distributions of response states rather than single deterministic values, information transmission was quantified at steady state as the mutual information between stimulus and downstream response distributions. Mutual information is a measure of dependence between two random variables and quantifies how much uncertainty about one variable (e.g. the stimulus) is reduced when the other variable (e.g. the response) is known [35]. In the context of cell signaling, mutual information measures how much information the response provides about the stimulus and therefore how reliably different stimulus concentrations can be discriminated [37, 10]. Analyses were restricted to stimulus concentrations producing between 10% and 90% or maximal receptor occupancy (see Methods and SI 1.8).

Notably, when DoRA approached 1 under normalization by 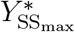, mutual information was minimal, indicating that the regime of maximal DoRA and near-linear occupancy–response behavior is associated with reduced information transmission (Fig. 3). In contrast, under normalization by *Y*_*T*_, the maxima of mutual information and DoRA occurred within similar *K*_*Y*_ */X*_*T*_ ranges, despite the occupancy–response relationship remaining far from linear (Fig. 3). Overall, these results show that neither perfect DoRA nor near-linear occupancy–response relationships necessarily maximize informa-tion transmission. Conversely, maximal information transmission can occur in regimes that remain non-linear.

**Figure 3:**
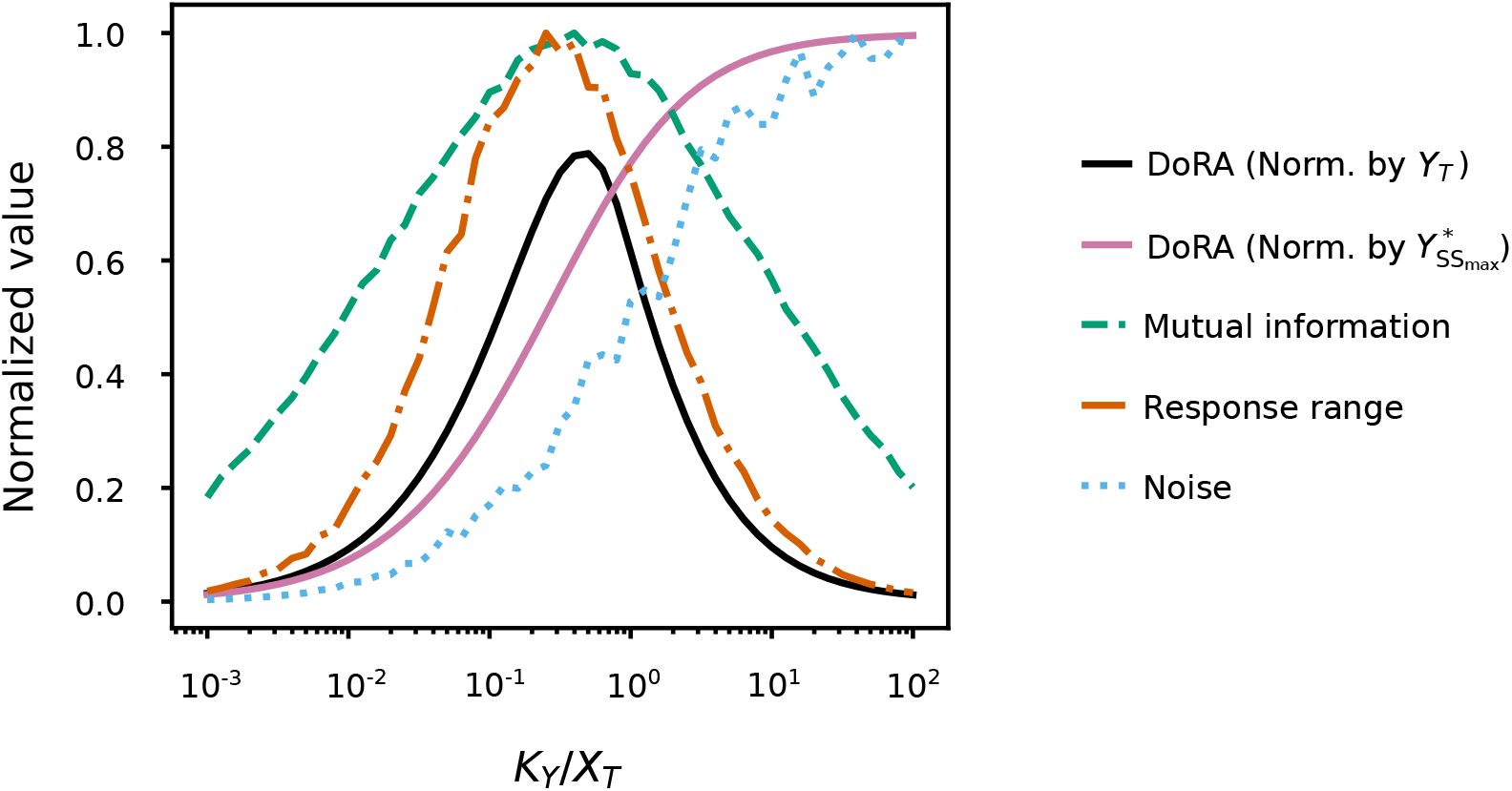
Relationship between DoRA, information transmission, response range, and noise. DoRA (under the two normalization schemes), mutual information, response range, and mean Fano factor are shown as functions of *K*_*Y*_ */X*_*T*_ . Under normalization by 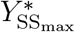, maximal DoRA occurs in a regime characterized by low mutual information, reduced response range, and elevated noise. Under normalization by *Y*_*T*_, maximal DoRA occurs near the regime of maximal mutual information, high response range, and intermediate noise levels. Mutual information, response range, and mean Fano factor were normalized to their respective maximum values.

Te decoupling between DoRA, occupancy–response linearity, and information transmission suggests that additional properties of the system may shape these relationships. Although DoRA quantifies the alignment between receptor occupancy and downstream responses, and occupancy–response linearity quantifies how closely the response follows a linear relationship with receptor occupancy, mutual information depends on the distinguishability of the full response distributions. This distinguishability is influenced by both the separation of the response distributions and their variability [28]. Response range contributes to information transmission by adjusting the separability of responses across stimuli [38, 28], whereas noise contributes by modulating the variability of the response distributions [39, 13, 28]. Moreover, we have shown that normalization alters the effective representation of response range and thereby affects the observed relationship between DoRA and occupancy–response linearity. These observations motivated us to examine the response range and noise as candidate properties underlying the observed relationship between DoRA and information transmission. Response range was defined as the difference between the maximum and minimum mean responses across stimulus concentrations, and noise was quantified as the Fano factor (variance-to-mean ratio of the response distribution) averaged over all stimulus conditions (see Methods and SI 1.9).

We found that, under normalization by 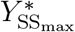, maximal DoRA occurred when the response range was minimal and noise maximal (Fig. 3). Under these conditions, the response distributions overlap strongly and exhibit high variability, consistent with reduced distinguishability across stimuli [40]. Consequently, mutual information is minimal (Fig. 3). In contrast, under normalization by *Y*_*T*_, maximal DoRA occurred alongside a broad response range and non-maximal noise levels (Fig. 3). These conditions increase the separation between response distributions and reduce their variability, enhancing their distinguishability. Importantly, mutual information was maximal at *K*_*Y*_ */X*_*T*_ values where response range remained high, and DoRA was close to its maximum (Fig. 3). Together, these results show that the relationship between DoRA and information transmission depends on how the normalization scheme shifts the position of maximal DoRA relative to linearity, response range, and noise, which jointly determine the distinguishability of response distributions (see SI 1.14).

## Discussion

In this study we investigated the relationship between dose–response alignment (DoRA), occupancy–response linearity and information transmission using deterministic and stochastic simulations of a simplified model of a cellular signaling pathway. Specifically we asked whether DoRA reliably captures occupancy-response linearity and whether it can serve as a proxy for the information transmission capacity of the system. Our re-sults show that these relationships are strongly influenced by the normalization scheme used to define downstream response. Consequently, high DoRA values do not necessarily reflect either linear occupancy–response relationship or enhanced information transmission. Together, these findings reveal that DoRA, linearity and information transmission, although related, represent distinct properties of signaling systems whose interpretation depends on how dose–response relationships are quantified.

DoRA is commonly associated with a correspondence between the range of receptor occupancies and the range of downstream responses [19]. However, it remains unclear whether the response range should be defined relative to the maximal response observed experimentally 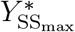 or relative to the total amount of response protein available in the system *Y*_*T*_ . Although both interpretations are consistent with the concept of DoRA, they imply different normalization schemes and therefore lead to distinct assessments of alignment. This ambiguity is not trivial, as experimental dose–response curves do not always allow one to determine whether the maximal observed response actually reaches the system’s full response capacity *Y*_*T*_, or whether it simply corresponds to the highest response attained under experimental conditions [41]. Our results show that this choice substantially affects the inferred degree of alignment: whereas normalization with respect to 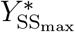 can yield perfect DoRA, normalization with respect to *Y*_*T*_ may result in considerably lower alignment values.

Our results further suggest that the magnitude of DoRA is not determined exclusively by the biochemical properties of the signaling system. For the same signaling pathway, with the same number of components and identical kinetic parameters, different normalization schemes can yield substantially different DoRA values. Consequently, the observed degree of alignment depends not only on the biochemical relationship between receptor occupancy and downstream response, but also on the representation used to define that response. In this sense, DoRA cannot be regarded as a purely intrinsic property of the signaling system, since it depends on how downstream responses are normalized and compared. These observations help reconcile why some studies report perfect DoRA in linear signaling pathways [23, 25], as is the case for our model. Such studies evaluate DoRA primarily in terms of proportionality, without considering whether the system reaches its full response capacity *Y*_*T*_ . Together, these findings highlight the inherent complexity of assessing DoRA [42].

We also found that the relationship between DoRA and occupancy–response linearity depends strongly on the normalization scheme. Occupancy–response linearity is often regarded as a desirable functional property of signaling pathways because it reflects the approximately proportional transmission of changes in receptor occupancy to downstream responses. In our model, this condition emerges when *K*_*Y*_ ≫ *X*_*T*_, where the downstream response becomes approximately proportional to receptor occupancy. Whereas normalization with respect to 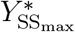 yielded near-perfect DoRA in this linear regime, normalization with respect to *Y*_*T*_ produced substantially lower DoRA values and shifted maximal alignment away from this linear regime. This difference arises because the two normalization schemes emphasize distinct system properties: normalization by 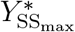 primarily captures proportionality between occupancy and response, whereas normalization by *Y*_*T*_ preserves the available dynamic range of the downstream response [24]. These results indicate that occupancy–response linearity and dynamic-range utilization are distinct properties of signaling systems, and that the extent to which DoRA reflects either one depends on how dose–response curves are normalized. This distinction becomes particularly important when considering information transmission, since dynamic range can influence the discriminability of responses independently of occupancy–response linearity.

Previous studies have suggested that DoRA is indicative of signaling systems with enhanced information transmission. The underlying intuition is that if the downstream response closely aligns with receptor occupancy (high DoRA), and if the occupancy– response relationship is nearly linear, then information should be transmitted more faithfully. Our results show that this is not necessarily the case. Although high DoRA under normalization by 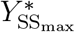 is associated with an approximately linear (and proportional) occupancy–response relationship, information transmission remains low. This occurs because linearity alone is insufficient to maximize information transmission if the response range is reduced. Under normalization by 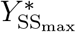, the occupancy–response relationship becomes nearly proportional, but the effective response range decreases substantially. As fewer response molecules become activated, relative noise becomes increasingly important, leading to greater overlap between response distributions and consequently lower mutual information.

The decoupling between DoRA and information transmission arises not only because DoRA does not uniquely capture occupancy–response linearity, depending on the normalization scheme employed, but also because DoRA and occupancy–response linearity are derived from mean dose–response relationships, whereas mutual information is a property of the full response distributions. In this sense, accurate information transmission depends not only on whether the mean downstream response aligns with receptor occupancy, but also on how response values are distributed around those means. From this perspective, DoRA primarily characterizes the geometry of normalized dose– response relationships, whereas information transmission depends on the separability of response distributions, which is jointly determined by response range and noise. Consequently, high alignment between occupancy and response does not necessarily imply enhanced information transmission if the corresponding response distributions remain poorly separated.

## Methods

### Model equations

We modeled the activation system as two coupled reversible reactions describing receptor occupancy and downstream response activation. The dynamic of these reactions were described by the following system of ordinary differential equations (ODEs):

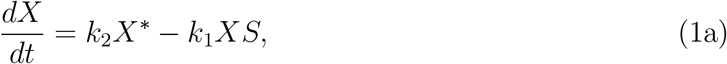

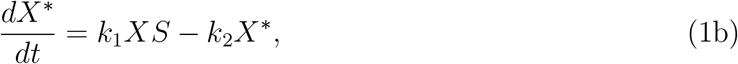

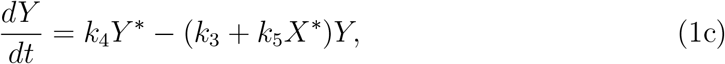

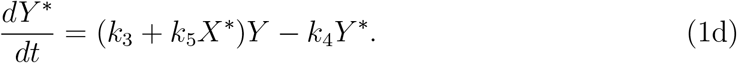

In these equations, *S* denotes the stimulus concentration, which was treated as an external input. The variables *X* and *X*\* represent the inactive and occupied receptor states, respectively, whereas *Y* and *Y* * denote the inactive and active forms of the response protein. The model assumes mass conservation such that *X*_*T*_ = *X* + *X*\* and *Y*_*T*_ = *Y* + *Y* *. The parameters *k*_1_ and *k*_2_ correspond to the association and dissociation rates between the receptor and the stimulus, respectively. The constants *k*_3_ and *k*_4_ describe the basal activation and inactivation rates of the response, whereas *k*_5_ quantifies the increase in response activation induced by occupied receptors (see SI 1.1).

### Steady states of the model

Setting the time derivatives to zero and assuming no basal activation of the response (*k*_3_ = 0; see SI 1.12 and 1.13 for the generalized case with *k*_3_ *>* 0), the steady-state solutions are:

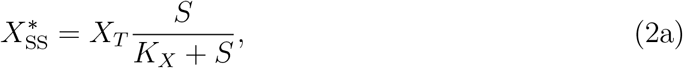

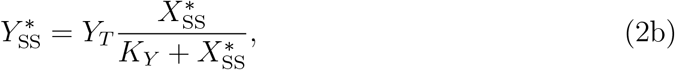

where *K*_*X*_ = *k*_2_*/k*_1_ and *K*_*Y*_ = *k*_4_*/k*_5_ correspond to the half-saturation constants for the receptor and downstream response, respectively (see SI 1.2).

### Normalized dose–response curves

Dose–response curves were obtained by evaluating the steady-state solutions over a logarithmically spaced stimulus range spanning three orders of magnitude above and below the receptor equilibrium constant *K*_*X*_ = *k*_2_*/k*_1_ (see SI 1.3). This ensured that the stimulus range was centered around the half-saturation receptor occupancy concentration.

We defined the normalized receptor occupancy and downstream response, respectively, as

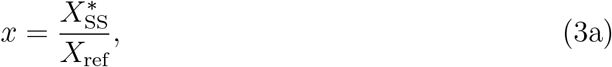

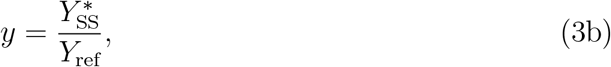

where *X*_ref_ and *Y*_ref_ denote the reference values used for normalization (see SI 1.4). Specifically, we considered 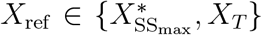 and 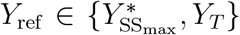, which correspond to the two normalization schemes described above.

### Dose-response alignment (DoRA) quantification

To quantify dose–response alignment (DoRA) between receptor occupancy and down-stream response curves, we used a generalized version of the metric introduced by [25] (see SI 1.5). This metric is defined as the weighted integral of the absolute differences between the normalized receptor occupancy and downstream response curves [25]:

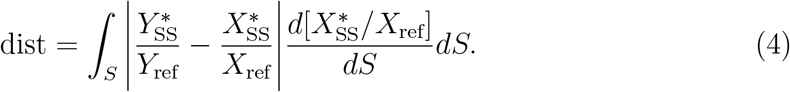

Because dist ranges from 0 to a maximal value of dist_max_ = 0.5, with zero corresponding to perfect alignment, we defined the DoRA measure as

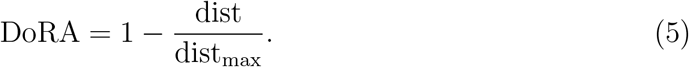

Values close to 1 indicate strong alignment, whereas values approaching 0 correspond to substantial misalignment.

As a complementary analysis, we also considered a transfer-function-based metric that yielded the same qualitative results; the corresponding analyses are presented in the SI 1.10 and 1.11.

### Linearity analysis

To evaluate occupancy–response linearity independently of normalization, we used the elasticity coefficient, a dimensionless measure from Metabolic Control Analysis that quantifies how relative changes in an input propagate to relative changes in an output [31, 32]. The elasticity coefficient is defined as

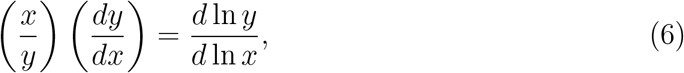

where a value of 1 indicates a locally linear relationship between input and output.

### Stochastic simulations

To estimate steady-state response probability distributions, we performed stochastic simulations of the model using the Gillespie stochastic simulation algorithm (SSA) [33, 34]. The algorithm explicitly accounts for intrinsic fluctuations arising from the discrete and stochastic nature of molecular interactions.

In the Gillespie framework, each reaction is assigned a propensity function that determines the relative probability of reaction events and the waiting time to the next event. The stochastic rate constants used in the simulations were derived from the corresponding deterministic rate constants as described in SI 1.7.

Because the Gillespie algorithm operates on molecular copy numbers rather than concentrations, all concentrations were converted into molecular copy numbers prior to simulation. To characterize steady-state fluctuations, simulations were initialized at the analytically calculated steady states (Eqs. (2a) and (2b)) and run sufficiently long to ensure adequate steady-state sampling. Specifically, each trajectory included at least 2,000 activation events of the downstream response species *Y* *. All stochastic simulations were implemented in Python 3.x using a custom Gillespie SSA code.

### Information quantification

Information transmission was quantified using the mutual information between the stimulus *S* and the downstream response *R*.

Here, *R* denotes the stochastic steady-state response corresponding to the copy number of the active response protein *Y* * (see SI 1.8). In this framework, *R* is treated as a discrete random variable representing the steady-state copy number of *Y* * obtained from stochastic simulations.

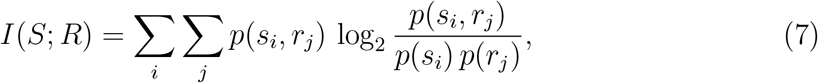

where *p*(*s*_*i*_, *r*_*j*_) denotes the joint probability distribution of stimulus and response values, and *p*(*s*_*i*_) and *p*(*r*_*j*_) are the corresponding marginal distributions.

Information was quantified within the transition region of the receptor dose–response curve, defined by the stimulus interval [*S*_10_, *S*_90_], corresponding to the stimulus concentrations producing 10% and 90% of maximal receptor occupancy, respectively. Stimulus concentrations were sampled uniformly on a logarithmic scale within this interval, generating a discrete set of stimulus values *s*_*i*_ used for the numerical evaluation of mutual information. This sampling strategy focuses on the transition region of the dose–response curve, where receptor occupancy changes substantially with stimulus concentration, while excluding regimes that contribute little to information transmission because the mean response varies minimally [6].

### Response range and noise estimation

The response range was quantified as the difference between the maximum and minimum mean steady-state responses across the stimulus range considered,

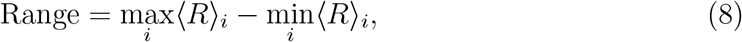

where ⟨*R*⟩_*i*_ denotes the mean response at stimulus concentration *s*_*i*_.

Intrinsic noise was quantified using the Fano factor,

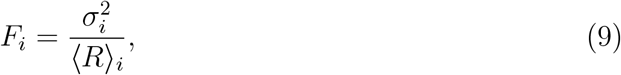

where σ^2^ is the variance of the response distribution at stimulus *s*_*i*_. The reported noise level corresponds to the average Fano factor across all stimulus concentrations considered. Details of the calculations are provided in SI 1.9.

## Acknowledgments

David Gordillo Alaniz is a doctoral student of the Programa de Doctorado en Ciencias Biomédicas at Universidad Nacional Autónoma de México (UNAM) and has received a fellowship (No. 1163254) from Secretaría de Ciencias, Humanidades, Tecnología e Innovación (SECIHTI, formerly CONAHCYT).

## References

[1] Bruce Alberts et al. Molecular Biology of the Cell. Sixth Edition. Garland Science, 2015.

[2] William Bialek and Sima Setayeshgar. “Physical limits to biochemical signaling”. In: PNAS 102 (29 2005), pp. 10040–10045.

[3] Jeremy E Purvis and Galit Lahav. “Encoding and decoding cellular information through signaling dynamics”. In: Cell 152 (5 2013), pp. 945–956.

[4] Andreas Wagner. “From bit to it: How a complex metabolic network transforms information into living matter”. In: BMC Syst. Biol. 1 (33 2007), pp. 1–9.

[5] Gašper Tkačik and William Bialek. “Information Processing in Living Systems”. In: Annu. Rev. Condens. Matter Phys. 7 (1 2016), pp. 89–117.

[6] Ryan Suderman and Eric J Deeds. “Intrinsic limits of information transmission in biochemical signalling motifs”. In: Interface Focus 8 (2018), p. 20180039.

[7] Alejandro Colman-Lerner et al. “Regulated cell-to-cell variation in a cell-fate decision system”. In: Nature 437 (7059 2005), pp. 699–706.

[8] Etay Ziv, Ilya Nemenman, and Chris H Wiggins. “Optimal signal processing in small stochastic biochemical networks”. In: PLoS ONE 2 (10 2007), e1077.

[9] John E Ladbury and Stefan T Arold. “Noise in cellular signaling pathways: Causes and effects”. In: Trends Biochem. Sci. 37 (5 May 2012), pp. 173–178.

[10] Clive G Bowsher and Peter S Swain. “Environmental sensing, information transfer, and cellular decision-making”. In: Curr. Opin. in Biotechnol. 28 (2014). Nanobiotechnology • Systems biology, pp. 149–155.

[11] Jonathan M Raser and Erin K O’shea. “Noise in Gene Expression: Origins, Consequences, and Control”. In: Science 309 (5743 Sept. 2025), pp. 2010–2013.

[12] Arjun Raj and Alexander van Oudenaarden. “Nature, Nurture, or Chance: Stochastic Gene Expression and Its Consequences”. In: Cell 135 (2 Oct. 2008), pp. 216–226.

[13] Alex Rhee, Raymond Cheong, and Andre Levchenko. “The application of information theory to biochemical signaling systems”. In: Phys. Biol. 9 (4 Aug. 2012), p. 045011.

[14] Piotr Topolewski and Micha-l Komorowski. “Information-theoretic analyses of cellular strategies for achieving high signaling capacity—dynamics, cross-wiring, and heterogeneity of cellular states”. In: Curr. Opin. Syst. Biol. 27 (2021), p. 100352.

[15] Gašper Tkačik and Pieter Rein Ten Wolde. “Information Processing in Biochemical Networks”. In: Annual Review ofBiophysics 52 (2025), p. 15.

[16] Diego A. Oyarzún et al. “The EGFR demonstrates linear signal transmission”. In: Integr. Biol. 6 (8 2014), pp. 736–742.

[17] Steven S Andrews, Roger Brent, and Gábor Balázsi. “Transferring information without distortion”. In: eLife 7 (2018), e41894.

[18] Harry Nunns and Lea Goentoro. “Signaling pathways as linear transmitters”. In: eLife 7 (2018), e33617.

[19] Richard C Yu et al. “Negative feedback that improves information transmission in yeast signalling”. In: Nature 456 (7223 2008), pp. 755–761.

[20] Margaritis Voliotis et al. “Information transfer by leaky, heterogeneous, protein kinase signaling systems”. In: PNAS 111 (3 2014), E326–E333.

[21] Louise Ashall et al. “Pulsatile Stimulation Determines Timing and Specificity of NF-kB-Dependent Transcription”. In: Science 324 (5924 Apr. 2009), pp. 242–246.

[22] Savas Tay et al. “Single-cell NF-B dynamics reveal digital activation and analogue information processing”. In: Nature 466 (7303 2010), pp. 267–271.

[23] Long Yan, Qi Ouyang, and Hongli Wang. “Dose-response aligned circuits in signaling systems”. In: PLoS ONE 7 (4 2012), e34727.

[24] Steven S Andrews et al. “Push-Pull and Feedback Mechanisms Can Align Signaling System Outputs with Inputs”. In: Cell Syst. 3 (5 2016), pp. 444–455.

[25] Lingxia Qiao, Pradipta Ghosh, and Padmini Rangamani. “Design principles of dose-response alignment in coupled GTPase switches”. In: npj Syst. Biol. Appl. 9 (3 2023).

[26] Roger Brent. “Cell signaling: What is the signal and what information does it carry?” In: FEBS Lett. 583 (24 2009), pp. 4019–4024.

[27] Marc Weimer et al. “The impact of data transformations on concentration-response modeling”. In: Toxicol. Lett. 213 (2 Sept. 2012), pp. 292–298.

[28] Eugenio Azpeitia, Eugenio P Balanzario, and Andreas Wagner. “Signaling pathways have an inherent need for noise to acquire information”. In: BMC Bioinform. 21 (462 2020), pp. 1–21.

[29] Anush Mahajan and Bhaswar Ghosh. “Optimization of Information Transmission through Noisy Biochemical Pathway”. In: Med. Res. Arch. 12 (3 2024), pp. 1–18.

[30] Khem Raj Ghusinga et al. “Molecular switch architecture determines response properties of signaling pathways”. In: PNAS 118 (11 2021), e2013401118.

[31] H Kacser and J Burns. “The control of flux”. In: Symp. Soc. Exp. Biol. 27 (1973), pp. 65–104.

[32] Reinhart Heinrich and Tom A Rapoport. “A Linear Steady-State Treatment of Enzymatic Chains: General Properties, Control and Effector Strength”. In: Eur. J. Biochem. 42 (1 1974), pp. 89–95.

[33] Daniel T Gillespie. “A General Method for Numerically Simulating the Stochastic Time Evolution of Coupled Chemical Reactions”. In: J. Comput. Phys. 22 (1976), pp. 403–434.

[34] Daniel T Gillespie. “Exact stochastic simulation of coupled chemical reactions”. In: J. Phys. Chem. 81 (25 1977), pp. 2340–2361.

[35] Thomas M Cover and Joy A Thomas. Elements of Information Theory. 2nd ed. Willey-Interscience, 2006, pp. 1–748.

[36] K A Zielińska and V L Katanaev. “Information Theory: New Look at Oncogenic Signaling Pathways”. In: Trends Cell Biol. 29 (11 2019), pp. 862–875.

[37] Filipe Tostevin and Pieter Rein ten Wolde. “Mutual information in time-varying biochemical systems”. In: Phys. Rev. E 81 (6 2010), p. 61917.

[38] Gašper Tkačik, Aleksandra M. Walczak, and William Bialek. “Optimizing information flow in small genetic networks”. In: Phys. Rev. E 80 (3 Sept. 2009).

[39] Raymond Cheong et al. “Information Transduction Capacity of Noisy Biochemical Signaling Networks”. In: Science 334 (6054 2011), pp. 354–358.

[40] Eugenio Azpeitia and Andreas Wagner. “Short Residence Times of DNA-Bound Transcription Factors Can Reduce Gene Expression Noise and Increase the Transmission of Information in a Gene Regulation System”. In: Front. Mol. Biosci. 7 (2020), p. 67.

[41] Terry P Kenakin. A Pharmacology Primer: Techniques for More Effective and Strategic Drug Discovery. 6th ed. Academic Press, 2022.

[42] John G Albeck, Michael Pargett, and Alexander E Davies. “Experimental and engineering approaches to intracellular communication”. In: Essays in Biochem. 62 (4 2018), pp. 515–524.

