## Supplementary Information for "Dose–Response Alignment Does Not Inherently Enhance Information Transmission in Signaling Pathways"

### 1 Supplementary Analyses

#### 1.1 System of ordinary differential equations (ODEs)

We model the signaling pathway using the following set of ordinary differential equations under the assumption of mass-action kinetics:

$$\frac{dX}{dt} = k_2X^* - k_1XS, \tag{1a}$$

$$\frac{dX^*}{dt} = k_1XS - k_2X^*, \tag{1b}$$

$$\frac{dY}{dt} = k_4Y^* - (k_3 + k_5X^*)Y, \tag{1c}$$

$$\frac{dY^*}{dt} = (k_3 + k_5X^*)Y - k_4Y^*. \tag{1d}$$

Here, $X$ and $X^*$ denote the concentrations for the inactive and active (occupied) forms of the receptor, respectively, while $Y$ and $Y^*$ represent the concentrations for the inactive and active forms of the response protein. The parameters $k_1$ and $k_2$ correspond to the association and dissociation rates between the receptor and the stimulus, respectively. The constants $k_3$ and $k_4$ describe the basal activation and inactivation rates of the response, and $k_5$ represents the receptor-mediated activation rate.

Using the conservation relations $X_T = X + X^*$ and $Y_T = Y + Y^*$, the system can be reduced to:

$$\frac{dX^*}{dt} = k_1(X_T - X^*)S - k_2X^*, \tag{2a}$$

$$\frac{dY^*}{dt} = (k_3 + k_5X^*)(Y_T - Y^*) - k_4Y^*. \tag{2b}$$

For the system to be well-defined and physically and biologically meaningful, we assume that the total concentrations ($X_T$ and $Y_T$) are strictly positive, while the state variables ($X^*$ and $Y^*$) and all the rate constants ($k_1, \ldots, k_5$) are non-negative.

#### 1.2 Steady states

Setting Eqs. (2a) and (2b) of the reduced system equal to zero yields the following steady-state expressions for $X^*$ and $Y^*$:

$$X^*_{\text{SS}} = X_T \frac{k_1S}{k_2 + k_1S}, \tag{3a}$$

$$Y^*_{\text{SS}} = Y_T \frac{k_3 + k_5X^*_{\text{SS}}}{k_3 + k_4 + k_5X^*_{\text{SS}}}. \tag{3b}$$

Although $Y^*_{\text{SS}}$ is written in terms of receptor activity, its dependence on the stimulus $S$ is fully determined through $X^*_{\text{SS}}$ (Eq. (3a)).

To isolate the effect of the receptor–response coupling, we focus on the scenario of zero basal activation ($k_3 = 0$). This assumption is physiologically reasonable, as many signaling pathways are expected to exhibit minimal activity levels in the absence of stimulus, ensuring that the response depends primarily on the stimulus [1]. With this assumption, the steady-state expressions (Eqs. (3a) and (3b)) can be rewritten as:

$$X^*_{\text{SS}} = X_T \frac{S}{K_X + S}, \tag{4a}$$

$$Y^*_{\text{SS}} = Y_T \frac{X^*_{\text{SS}}}{K_Y + X^*_{\text{SS}}}. \tag{4b}$$

Both expressions follow a Michaelis–Menten saturation kinetics, in which $K_X = k_2/k_1$ and $K_Y = k_4/k_5$ determine the stimulus concentration required to reach half-maximal receptor and downstream response activation, respectively. In other words, $K_X$ and $K_Y$ act as effective half-saturation constants that determine the shape of the receptor occupancy and downstream curves, respectively.

#### 1.3 Dose–response curves

Dose–response curves relate the dose of an active agent, such as a ligand, to its effect in a biological system. In practice, these curves are typically defined in terms of the steady-state response associated with a given dose [2]. In our model, these curves are given by the steady-state levels of the occupied receptor, $X^*_{\mathrm{SS}}$, and the active response protein, $Y^*_{\mathrm{SS}}$, as a function of the stimulus $S$.

To construct the dose–response curves, the steady-state expressions $X^*_{\mathrm{SS}}$ and $Y^*_{\mathrm{SS}}$ were evaluated using Eqs. (4a) and (4b) over 100 logarithmically spaced stimulus concentrations ranging from $10^{-3}K_X$ to $10^{3}K_X$, where $K_X = k_2/k_1$. The resulting curves were subsequently normalized to the procedure described in the following section.

#### 1.4 Normalization of dose–response curves

Since dose–response curves of receptor occupancy and downstream responses typically differ in magnitude, they are commonly normalized to a common scale to comparison [3].

In general, a response $R$ can be normalized by considering its minimum ($R_{\mathrm{min}}$) and maximum ($R_{\mathrm{max}}$) values as

$$R_{\mathrm{norm}} = \frac{R - R_{\mathrm{min}}}{R_{\mathrm{max}} - R_{\mathrm{min}}}, \tag{5}$$

where $R_{\mathrm{min}}$ corresponds to the value of the response in the absence of the stimulus, while $R_{\mathrm{max}}$ denotes the maximum response value. However, experimentally is not always clear whether the $R_{\mathrm{max}}$ is equal to the full capacity of the system or merely to the maximum reached under specific conditions considered [4].

In our model, this normalization is applied to the steady-state responses $X^*_{\mathrm{SS}}$ and $Y^*_{\mathrm{SS}}$. We define the dimensionless variables $x$ and $y$, representing the normalized receptor occupancy and downstream responses, respectively, as

$$x = \frac{X^*_{\mathrm{SS}} - X^*_{\mathrm{SS_{min}}}}{X_{\mathrm{ref}} - X^*_{\mathrm{SS_{min}}}}, \tag{6a}$$

$$y = \frac{Y^*_{\mathrm{SS}} - Y^*_{\mathrm{SS_{min}}}}{Y_{\mathrm{ref}} - Y^*_{\mathrm{SS_{min}}}}, \tag{6b}$$

where $X_{\mathrm{ref}}$ and $Y_{\mathrm{ref}}$ are reference values that determine the normalization scale. In this work, they can correspond either to the maximum steady-state responses reached under the conditions considered ($X^*_{\mathrm{SS_{max}}}$, $Y^*_{\mathrm{SS_{max}}}$), or to the total concentrations of the receptor and response protein ($X_T$, $Y_T$), thereby defining different normalization schemes.

Under zero basal activity ($k_3 = 0$), the previous normalized responses (Eqs. (6a) and (6b)) reduce to:

$$x = \frac{X_{\mathrm{SS}}^*}{X_{\mathrm{ref}}}, \tag{7}$$

$$y = \frac{Y_{\mathrm{SS}}^*}{Y_{\mathrm{ref}}}. \tag{8}$$

Naturally, when $X^*_{\mathrm{SS_{max}}} = X_T$ and $Y^*_{\mathrm{SS_{max}}} = Y_T$, both normalization schemes are equivalent. Otherwise, the resulting normalized dose–response curves differ because they are scaled using different reference values.

The normalized dose–response curves are defined as the normalized responses $x$ and $y$, as a function of the stimulus $S$.

Thus, normalization is not merely a rescaling procedure; it can modify the relative shape of the curves and therefore influence the quantification of dose–response alignment and the information transmitted by the system.

#### 1.5 Dose–response alignment (DoRA)

By eliminating the explicit dependence of the stimulus $S$, the normalized dose–response curves define an occupation–response relation of the form

$$y = f(x), \tag{9}$$

where $x$ and $y$ denote the normalized receptor occupancy and downstream response curves, respectively. This relationship, often referred to as the *transfer function* of the system, provides a geometric description of how normalized receptor occupancy is mapped onto normalized downstream response, independently of the original physical units or the stimulus range.

Within this framework, perfect dose–response alignment (DoRA) corresponds to the identity mapping,

$$f(x) = x, \tag{10}$$

or equivalently,

$$f(x) - x = 0, \tag{11}$$

for all $x \in [0,1]$, since both responses are normalized. Any deviation from this equality reflects misalignment between normalized receptor occupancy and normalized downstream response.

A natural way to quantify this deviation is through the area between the transfer function and the identity line, which defines a distance measure between the two curves,

$$\text{dist} = \int_0^1 |f(x) - x| dx. \tag{12}$$

In practice, this distance measure can be evaluated directly from the steady-state dose–response curves parametrized by the stimulus $S$. Applying the change of variable $dx = (dx/dS)dS$, the distance becomes

$$\text{dist} = \int_S |y(S) - x(S)| \frac{dx}{dS} dS. \tag{13}$$

In the scenario of zero basal activity ($k_3 = 0$), the normalized responses correspond to Eqs (7) and (8), yielding

$$\text{dist} = \int_S \left| \frac{Y^*_{\text{SS}}}{Y_{\text{ref}}} - \frac{X^*_{\text{SS}}}{X_{\text{ref}}} \right| \frac{d\left(X^*_{\text{SS}}/X_{\text{ref}}\right)}{dS} dS. \tag{14}$$

When $X_{\text{ref}} = X^*_{\text{SS}_{\text{max}}}$ and $Y_{\text{ref}} = Y^*_{\text{SS}_{\text{max}}}$, this expression reduces to the formulation of [5].

Finally, since the distance is bounded between 0 and a maximal value $\text{dist}_{\text{max}} = 1/2$, we define a normalized DoRA measure as

$$\text{DoRA} = 1 - \frac{\text{dist}}{\text{dist}_{\text{max}}}, \tag{15}$$

which assigns a value of 1 to perfect alignment and a value of 0 to maximal misalignment.

#### 1.6 Linearity and the elasticity coefficient

To evaluate whether the relationship between receptor occupancy and response is linear independently of normalization, we used the elasticity coefficient, a dimensionless measure that quantifies how relative changes in an input are transmitted as relative changes in an output. This quantity has been widely used in Metabolic Control Analysis [6, 7]. In particular, when the elasticity coefficient is equal to 1, the relationship between input and output is locally linear. The elasticity coefficient is defined as:

$$\left(\frac{x}{y}\right)\left(\frac{dy}{dx}\right) = \frac{d\ln y}{d\ln x} \tag{16}$$

For our model, we evaluated the change in $Y^*_{\text{SS}}$ as a function of $X^*_{\text{SS}}$, such that the elasticity coefficient is given by:

$$\left(\frac{X^*_{\text{SS}}}{Y^*_{\text{SS}}}\right)\left(\frac{dY^*_{\text{SS}}}{dX^*_{\text{SS}}}\right) = \left(\frac{K_Y + X^*_{\text{SS}}}{Y_T}\right)\left(\frac{Y_T K_Y}{(K_Y + X^*_{\text{SS}})^2}\right) = \frac{K_Y}{K_Y + X^*_{\text{SS}}} \tag{17}$$

From this expression, it can be observed that because $X^*_{\text{SS}}$ is upper bounded by $X_T$, when the ratio $K_Y/X_T \to \infty$, it follows that $X^*_{\text{SS}}/K_Y \to 0$ for all admissible values of $X^*_{\text{SS}}$. Consequently, the elasticity coefficient approaches 1 throughout the full range of $X^*_{\text{SS}}$:

$$\frac{K_Y}{K_Y + X^*_{\text{SS}}} = \frac{1}{1 + X^*_{\text{SS}}/K_Y} \to 1$$

Thus, although the elasticity coefficient is formally a local measure, under the condition $K_Y/X_T \to \infty$ the occupancy–response relationship becomes arbitrarily close to linear across its entire dynamic range.

For the general case $k_3 \geq 0$, the elasticity coefficient of the downstream response with respect to receptor activation is

$$\frac{d\ln y}{d\ln x} = \frac{k_4 k_5 X^*_{\text{SS}}}{(k_3 + k_5 X^*_{\text{SS}})\,(k_3 + k_4 + k_5 X^*_{\text{SS}})}.$$

When $k_3 > 0$, the denominator can be rewritten as

$$(k_3 + k_5 X^*_{\text{SS}})\,(k_3 + k_4 + k_5 X^*_{\text{SS}}) = k_3(k_3 + k_4 + 2k_5 X^*_{\text{SS}}) + (k_5 X^*_{\text{SS}})^2 + k_4 k_5 X^*_{\text{SS}}.$$

Since all terms are strictly positive for $k_3 > 0$,

$$k_3(k_3 + k_4 + 2k_5 X^*_{\text{SS}}) + (k_5 X^*_{\text{SS}})^2 + k_4 k_5 X^*_{\text{SS}} > k_4 k_5 X^*_{\text{SS}},$$

and therefore

$$\frac{d\ln y}{d\ln x} < 1.$$

Consequently, for any positive basal activation rate $k_3 > 0$, the downstream module is intrinsically sublinear. Furthermore, because $k_3$ appears only in the denominator, increasing basal receptor activation decreases the elasticity coefficient and thus enhances the degree of sublinearity.

Nevertheless, in the asymptotic regime $k_4 \gg k_3$, the elasticity can be approximated as

$$\frac{d\ln y}{d\ln x} \approx \frac{k_4 k_5 X^*_{\mathrm{SS}}}{(k_3 + k_5 X^*_{\mathrm{SS}})\,(k_4 + k_5 X^*_{\mathrm{SS}})}.$$

If, in addition,

$$k_5 X^*_{\mathrm{SS}} \gg k_3,$$

the contribution of basal receptor activation becomes negligible, yielding

$$\frac{d\ln y}{d\ln x} \approx \frac{k_4}{k_4 + k_5 X^*_{\mathrm{SS}}}.$$

Using $K_Y = k_4/k_5$,

$$\frac{d\ln y}{d\ln x} \approx \frac{K_Y}{K_Y + X^*_{\mathrm{SS}}} = \frac{1}{1 + X^*_{\mathrm{SS}}/K_Y}.$$

Therefore, when

$$k_3 \ll k_5 X^*_{\mathrm{SS}} \ll k_4,$$

the elasticity asymptotically reduces to the same expression obtained for $k_3 = 0$. Consequently,

$$\frac{K_Y}{X^*_{\mathrm{SS}}} \to \infty \quad \Longrightarrow \quad \frac{d\ln y}{d\ln x} \to 1.$$

Since $X^*_{\mathrm{SS}} \leq X_T$, the stronger condition

$$\frac{K_Y}{X_T} \to \infty$$

is sufficient to ensure arbitrarily close-to-unit elasticity across the full admissible range of receptor activation.

#### 1.7 Stochastic simulations

To estimate the response probability distributions, we performed stochastic simulations of the model using the Gillespie stochastic simulation algorithm (SSA) [8, 9].

The Gillespie formalism considers a homogeneous system composed of $N$ chemical species $Z_1, \ldots, Z_N$, interacting through $M$ chemical reactions $\mathcal{R}_1, \ldots, \mathcal{R}_M$. The state of the system at time $t$ is represented by the vector

$$n(t) = (n_1(t), \ldots, n_N(t)),$$

where $n_i(t)$ denotes the number of molecules of species $Z_i$.

The Gillespie algorithm simulates the temporal evolution of the system by determining both the waiting time $\tau$ until the next reaction event and the reaction $\mathcal{R}_\mu$ that occurs. To this end, each reaction $\mathcal{R}_\mu$ is characterized by a propensity function

$$a_\mu = h_\mu c_\mu,$$

where $h_\mu$ is the number of distinct reactant combinations available for reaction $\mathcal{R}_\mu$, and $c_\mu$ is the stochastic rate constant associated with that reaction. The stochastic rate constants can be derived from the corresponding deterministic rate constants $k_\mu$ according to the reaction order, as summarized in Table S1.

Table S1: Stochastic parameters for each type of reaction

| Reaction order | Reaction | $h$ | $c$ |
|---|---|---|---|
| Zero-order | $\emptyset \xrightarrow{k} Z_1$ | 1 | $kVN_A$ |
| First-order | $Z_1 \xrightarrow{k} Z_2$ | $Z_1$ | $k$ |
| Second-order | $Z_1 + Z_2 \xrightarrow{k} Z_3$ | $Z_1 Z_2$ | $\frac{k}{VN_A}$ |
| Second-order | $Z_1 + Z_1 \xrightarrow{k} Z_2$ | $\frac{Z_1(Z_1-1)}{2}$ | $\frac{k}{VN_A}$ |

Since the Gillespie algorithm operates on molecule counts, concentrations for all variables were converted into molecular copy numbers using the definition of molar concentration:

$$n_i = [Z_i]\, V\, N_A \tag{18}$$

where $n_i$ is the number of molecules of species $Z_i$, $[Z_i]$ is the molar concentration of $Z_i$, $V$ is the system volume, and $N_A$ is Avogadro's number.

The reactions involved in the model are

$$
\begin{aligned}
\mathcal{R}_1 &: X + S \xrightarrow{k_1} X^*, \\
\mathcal{R}_2 &: X^* \xrightarrow{k_2} X + S, \\
\mathcal{R}_3 &: Y \xrightarrow{k_3} Y^*, \\
\mathcal{R}_4 &: Y^* \xrightarrow{k_4} Y, \\
\mathcal{R}_5 &: X^* + Y \xrightarrow{k_5} Y^* + X^*.
\end{aligned}
$$

Using Table S1 and Eq. (18), the propensities associated with reactions $\mathcal{R}_1$ to $\mathcal{R}_5$ are

$$
\begin{aligned}
a_1 &= n_X n_S \frac{k_1}{N_A V}, \\
a_2 &= n_{X^*} k_2, \\
a_3 &= n_Y k_3, \\
a_4 &= n_{Y^*} k_4, \\
a_5 &= n_{X^*} n_Y \frac{k_5}{N_A V}.
\end{aligned}
$$

Note that for the scenario of zero basal response ($k_3 = 0$), the reaction $\mathcal{R}_3$ and its associated propensity $a_3$ do not occur.

Because we are interested in the steady-state probability distribution of the response, simulations were initialized at the analytically calculated steady state (Eqs. (4a) and 4b). Since the system satisfies the ergodic property, a single sufficiently long stochastic trajectory was sufficient to estimate the steady-state distribution. The simulation length was chosen to ensure that the stochastic trajectory included at least 2,000 reaction events involving the activation of the downstream response ($Y^*$), which guaranteed adequate sampling of the steady-state fluctuations.

#### 1.8 Information

Mutual information is a measure of dependence between two random variables and quantifies how much uncertainty about one variable is reduced due to the knowledge of the other variable [10]. In biochemical signaling systems, it has become an important measure of information transmission [11, 12]. Here, we used mutual information to quantify information transmission between the stimulus $S$ and the downstream response $R$. In our model, $R$ corresponds to the stochastic abundance of the active response protein $Y^*$ at steady state.

The mutual information between the stimulus and the response is defined as

$$I(S;R) = \sum_i \sum_j p(s_i, r_j) \, \log_2 \frac{p(s_i, r_j)}{p(s_i) \, p(r_j)}, \tag{19}$$

where $p(s_i, r_j)$ is the joint probability distribution of the stimulus and response values, and $p(s_i)$ and $p(r_j)$ are the corresponding marginal probability distributions. If $S$ and $R$ are statistically independent, then $I(S;R) = 0$, indicating that the response carries no information about the stimulus. Conversely, if $S$ and $R$ are identical, so that knowing the value of one determines the value of the other, then the stimulus is transmitted perfectly to the response, and the mutual information reaches its maximum possible value for those variables.

As discussed by [13], most of the information transmitted through a sigmoidal dose–response curve is concentrated within what they term the *increasing regime* or the *transition zone*, typically spanning the stimulus interval $[S_{10}, S_{90}]$, corresponding to the stimulus concentrations producing 10% and 90% of the maximal response, respectively. Accordingly, we quantified mutual information using $n$ stimulus concentrations uniformly distributed on a logarithmic scale within the interval $[S_{10}, S_{90}]$ of the receptor occupancy dose–response curve, as receptor occupancy defines the maximal information that can be transmitted downstream through the signaling pathway.

Since the system satisfies the ergodic property, steady-state response distributions could be estimated from a single sufficiently long stochastic trajectory for each stimulus concentration $s_i$ (see SI 1.7). The conditional response distribution $p(r_j|s_i)$ was therefore estimated as the fraction of simulation time that the system spent in each response state $r_j$. Here, $r_j$ denotes a possible steady-state abundance of the active response protein $Y^*$.

Stimulus concentrations were assumed to be equiprobable, such that

$$p(s_i) = \frac{1}{n}. \tag{20}$$

The marginal response distribution was then obtained as

$$p(r_j) = \sum_{i=1}^{n} p(r_j \mid s_i) \, p(s_i), \tag{21}$$

and the joint distribution was computed according to

$$p(s_i, r_j) = p(r_j \mid s_i) \, p(s_i). \tag{22}$$

#### 1.9 Response range and noise

The response range quantifies the extent of variation in the average downstream response over the stimulus interval considered (see SI 1.8). Here, $R$ corresponds to the stochastic abundance of the active response protein $Y^*$ at steady state. For each stimulus concentration $s_i$, the mean response was obtained from the conditional probability distribution $p(r_j \mid s_i)$ as

$$\langle R \rangle_i = \sum_{r_j} r_j \, p(r_j \mid s_i). \tag{23}$$

The response range was then defined as the difference between the maximum and the minimum mean responses across all stimulus concentrations,

$$\text{Range} = \max_i \langle R \rangle_i - \min_i \langle R \rangle_i. \tag{24}$$

where the maximum and minimum are taken over all stimulus concentrations considered.

The level of intrinsic noise in the downstream response was quantified using the Fano factor, defined as the ratio between the variance and the mean response. For each stimulus concentration $s_i$, the variance of the response distribution was computed as

$$\sigma_i^2 = \sum_{r_j} (r_j - \langle R \rangle_i)^2 \, p(r_j \mid s_i). \tag{25}$$

The Fano factor associated with stimulus $s_i$ was calculated as

$$F_i = \frac{\sigma_i^2}{\langle R \rangle_i}. \tag{26}$$

Finally, the average Fano factor was obtained by averaging over all stimulus concentrations considered,

$$\overline{F} = \frac{1}{n} \sum_{i=1}^{n} F_i. \tag{27}$$

#### 1.10 Dose–response alignment when normalized by maximum responses

First we considered the maximum observed responses as reference values (i.e., $X_{\text{ref}} = X^*_{\text{SS}_{\max}}$ and $Y_{\text{ref}} = Y^*_{\text{SS}_{\max}}$) to construct normalized dose–response curves. Under this scheme, the Eqs. (6a) and (6b) become:

$$x = \frac{X^*_{\mathrm{SS}} - X^*_{\mathrm{SS_{min}}}}{X^*_{\mathrm{SS_{max}}} - X^*_{\mathrm{SS_{min}}}}, \tag{28a}$$

$$y = \frac{Y^*_{\mathrm{SS}} - Y^*_{\mathrm{SS_{min}}}}{Y^*_{\mathrm{SS_{max}}} - Y^*_{\mathrm{SS_{min}}}}. \tag{28b}$$

Since we consider the case of zero basal activity ($k_3 = 0$), the steady-state expressions for receptor occupancy and downstream responses are given by Eqs. (4a) and (4b):

$$X^*_{\mathrm{SS}} = X_T \frac{S}{K_X + S}, \tag{29}$$

$$Y^*_{\mathrm{SS}} = Y_T \frac{X^*_{\mathrm{SS}}}{K_Y + X^*_{\mathrm{SS}}}, \tag{30}$$

From these expressions, the minimum and maximum response values can be directly obtained. For the receptor, the minimum occurs when in the absence of stimulus ($S \to 0$), while the maximum is reached as $S \to \infty$, yielding:

$$X^*_{\mathrm{SS_{min}}} = 0, \tag{31}$$

$$X^*_{\mathrm{SS_{max}}} = X_T. \tag{32}$$

For the downstream response, the minimum and maximum values correspond to the minimum and maximum receptor occupancy, respectively. Therefore:

$$Y^*_{\mathrm{SS_{min}}} = 0, \tag{33}$$

$$Y^*_{\mathrm{SS_{max}}} = Y_T \frac{X_T}{K_Y + X_T}. \tag{34}$$

Substituting these expressions into Eqs. (28a) and (28b), we obtain the normalized dose–response curves:

$$x = \frac{X^*_{\mathrm{SS}}}{X^*_{\mathrm{SS_{max}}}} = \frac{X^*_{\mathrm{SS}}}{X_T}, \tag{35a}$$

$$y = \frac{Y^*_{\mathrm{SS}}}{Y^*_{\mathrm{SS_{max}}}} = \frac{X^*_{\mathrm{SS}}}{X_T} \frac{K_Y + X_T}{K_Y + X^*_{\mathrm{SS}}}. \tag{35b}$$

Rewriting $x$ as $X^*_{\text{SS}} = X_T x$ and substituting into $y$, we obtain the transfer function:

$$f(x) = \frac{x(K_Y + X_T)}{K_Y + X_T x}. \tag{36}$$

To determine whether perfect dose–response alignment (DoRA) can be achieved under this normalization scheme, we calculated the difference between the curves by substituting Eq. (36) into Eq. (11):

$$f(x) - x = \frac{X_T x(1-x)}{K_Y + X_T x}, \tag{37}$$

Perfect alignment requires that the above equation be equal to zero for all $x \in [0, 1]$. Since $X_T > 0$ and $K_Y > 0$, the denominator is strictly positive over this interval. Therefore, the condition can only be satisfied when the numerator is zero, which occurs at $x = 0$ and $x = 1$, corresponding respectively to zero receptor occupancy and full saturation.

Thus, the two curves coincide only at the boundaries of the interval. For any $0 < x < 1$, we have $f(x) \neq x$, indicating that perfect dose–response alignment (DoRA) cannot be achieved under this normalization scheme.

To gain further insight, we rewrite Eq. (37) in terms of the ratio $K_Y/X_T$ as

$$f(x) - x = \frac{x(1-x)}{\frac{K_Y}{X_T} + x}.$$

From this expression, it follows that in the limit $K_Y/X_T \to \infty$ (or equivalently $X_T/K_Y \to 0$), the deviation between the two curves approaches zero:

$$\lim_{\frac{K_Y}{X_T} \to \infty} (f(x) - x) = 0,$$

and therefore the transfer function $f(x)$ approaches the identity mapping $f(x) = x$.

Thus, although perfect DoRA cannot be achieved for finite parameter values, it emerges asymptotically in the limit $K_Y/X_T \to \infty$, corresponding to regimes in which $K_Y$ becomes much larger than $X_T$.

#### 1.11 Dose–response alignment when normalized by total protein available response protein

When normalization is performed with respect to the total amounts of protein (i.e., $X_{\text{ref}} = X_T$ and $Y_{\text{ref}} = Y_T$) to construct normalized dose–response curves, Eqs. (6a) and (6b) become

$$x = \frac{X^*_{\text{SS}} - X^*_{\text{SS}_{\text{min}}}}{X_T - X^*_{\text{SS}_{\text{min}}}}, \tag{38a}$$

$$y = \frac{Y^*_{\text{SS}} - Y^*_{\text{SS}_{\text{min}}}}{Y_T - Y^*_{\text{SS}_{\text{min}}}}. \tag{38b}$$

Under zero basal activity ($k_3 = 0$), the minimum steady-state responses remain identical to those obtained in the previous normalization scheme, namely $X^*_{\text{SS}_{\text{min}}} = 0$ and $Y^*_{\text{SS}_{\text{min}}} = 0$. Under this normalization scheme, the reference values are given by the total amounts of protein $X_T$ and $Y_T$. Substituting these conditions into Eqs. (38a) and (38b) yields

$$x = \frac{X^*_{\text{SS}}}{X_T}, \tag{39a}$$

$$y = \frac{Y^*_{\text{SS}}}{Y_T} = \frac{X^*_{\text{SS}}}{K_Y + X^*_{\text{SS}}}. \tag{39b}$$

Note that the normalized receptor occupancy obtained under this scheme (Eq. 39a) is identical to that obtained when normalization is performed with respect to the maximum response (Eq. 35a), since for the receptor $X^*_{\text{SS}_{\text{max}}} = X_T$. Thus, receptor occupancy is always expressed as a fraction of the total receptor available.

The downstream response curve, however, differs between normalization schemes. Rewriting Eq. (39a) as $X^*_{\text{SS}} = X_T x$ and substituting into Eq. (39b), we obtain the corresponding transfer function

$$f(x) = \frac{X_T x}{K_Y + X_T x}. \tag{40}$$

To evaluate whether perfect dose–response alignment (DoRA) can occur under this normalization scheme, we computed the deviation from the identity mapping by substituting Eq. (40) into Eq. (11):

$$f(x) - x = \frac{x(X_T(1-x) - K_Y)}{K_Y + X_T x}. \tag{41}$$

Perfect alignment requires $f(x) = x$ for all $x \in [0, 1]$. Since the denominator remains strictly positive for $K_Y > 0$, the intersections between the two curves are determined by the numerator. The curves always intersect at $x = 0$, corresponding to zero receptor occupancy. A second intersection occurs when

$$X_T(1 - x) - K_Y = 0, \tag{42}$$

which gives

$$x = 1 - \frac{K_Y}{X_T}. \tag{43}$$

This second solution lies within the normalized activation interval only when $K_Y \leq X_T$; otherwise, the two curves intersect exclusively at $x = 0$. Thus, unlike the normalization scheme based on maximal response, the two curves do not coincide at both boundaries of the activation interval, and perfect DoRA cannot be achieved across the full activation range $0 \leq x \leq 1$.

For convenience, the transfer function can also be written as

$$f(x) = \frac{x}{\frac{K_Y}{X_T} + x}. \tag{44}$$

In contrast to the previous normalization scheme, this expression does not approach the identity mapping $f(x) = x$ in any asymptotic parameter regime. Thus, perfect alignment does not emerge even asymptotically.

So, although perfect alignment cannot be achieved under normalization by total protein abundance, the deviation between the receptor occupancy and downstream response curves still depends on the ratio $K_Y/X_T$. This raises the question of which ratio yields the highest degree of dose–response alignment.

To address this question, we used the same distance measure introduced in Eq. (12). Using the transfer function in Eq. (44), the distance becomes

$$D\left(\frac{K_Y}{X_T}\right) = \int_0^1 \left| \frac{x}{\frac{K_Y}{X_T} + x} - x \right| dx, \tag{45}$$

Since DoRA is directly determined by the distance between the receptor occupancy and downstream response curves (Eq. 15), the value of the ratio $K_Y/X_T$ that maximizes DoRA correspond to the value that minimizes $D(K_Y/X_T)$. Numerical evaluation of $D(K_Y/X_T)$ yields an optimal ratio of

$$\frac{K_Y}{X_T} \approx 0.4622.$$

Thus, maximal dose–response alignment under normalization with respect to total protein abundance $Y_T$ emerges at an intermediate ratio $K_Y/X_T$.

#### 1.12 Dose–response alignment when normalized by maximum observed response for $k_3 > 0$

We now examine the case when the basal rate $k_3 > 0$. In this scenario the steady-state expression for the downstream response is given by Eq. (3b)

$$Y^*_{\mathrm{SS}} = Y_T \frac{k_3 + k_5 X^*_{\mathrm{SS}}}{k_3 + k_4 + k_5 X^*_{\mathrm{SS}}},$$

and dividing both the numerator and denominator by $k_5$ we obtain

$$Y^*_{\mathrm{SS}} = Y_T \frac{X^*_{\mathrm{SS}} + \frac{k_3}{k_5}}{K_Y + X^*_{\mathrm{SS}} + \frac{k_3}{k_5}}. \tag{46}$$

where $K_Y = k_4/k_5$ is the half-saturation constant for the downstream response.

Since the receptor dynamics do not depend on the rate $k_3$, its minimum and maximum responses remain identical to those obtained in the previous case (Eqs. (31) and 32),

$$X^*_{\mathrm{SS}_{\min}} = 0,$$
$$X^*_{\mathrm{SS}_{\max}} = X_T.$$

Likewise, the maximum downstream response is reached when $X^*_{\mathrm{SS}} \to X_T$. Evaluating Eq. (46) in this limit gives

$$Y^*_{\mathrm{SS}_{\max}} = Y_T \frac{X_T + \frac{k_3}{k_5}}{K_Y + X_T + \frac{k_3}{k_5}}.$$

The minimum downstream response is also modified by the presence of basal activity. In this case, the response protein remains at a nonzero activation level even when $X^*_{\mathrm{SS}} \to 0$. Evaluating this limit in the above steady-state expression (Eq. (46)) yields

$$Y^*_{\mathrm{SS}_{\min}} = Y_T \frac{X^*_{\mathrm{SS}_{\min}} + \frac{k_3}{k_5}}{K_Y + X^*_{\mathrm{SS}_{\min}} + \frac{k_3}{k_5}} = Y_T \frac{\frac{k_3}{k_5}}{K_Y + \frac{k_3}{k_5}}. \tag{47}$$

By substituting the minimum and maximum response values in Eq. (28b) we obtain the following expressions for the normalized dose–response curve:

$$y = \frac{Y^*_{\mathrm{SS}} - Y^*_{\mathrm{SS_{min}}}}{Y^*_{\mathrm{SS_{max}}} - Y^*_{\mathrm{SS_{min}}}} = \frac{Y_T\left[\frac{X^*_{\mathrm{SS}}+\frac{k_3}{k_5}}{K_Y+X^*_{\mathrm{SS}}+\frac{k_3}{k_5}} - \frac{\frac{k_3}{k_5}}{K_Y+\frac{k_3}{k_5}}\right]}{Y_T\left[\frac{X_T+\frac{k_3}{k_5}}{K_Y+X_T+\frac{k_3}{k_5}} - \frac{\frac{k_3}{k_5}}{K_Y+\frac{k_3}{k_5}}\right]}, \tag{48}$$

which can be simplified as

$$y = \frac{X^*_{\mathrm{SS}}(K_Y + X_T + \frac{k_3}{k_5})}{X_T(K_Y + X^*_{\mathrm{SS}} + \frac{k_3}{k_5})}. \tag{49}$$

We now define $f(x) = y$ from the expression above and substitute $X^*_{\mathrm{SS}} = X_T x$, obtaining

$$f(x) = \frac{x\left(K_Y + X_T + \frac{k_3}{k_5}\right)}{K_Y + X_T x + \frac{k_3}{k_5}}, \tag{50}$$

and from it we calculate the deviation from the identity mapping as

$$f(x) - x = \frac{X_T x(1-x)}{K_Y + X_T x + \frac{k_3}{k_5}}. \tag{51}$$

This expression has the same structure as the corresponding result obtained for $k_3 = 0$ (Eq. 37). Consequently, as in the case $k_3 = 0$, the system fails to produce perfect dose–response alignment. The two curves coincide only at receptor inactivity ($x = 0$) and receptor saturation ($x = 1$). For all $0 < x < 1$ , the normalized downstream response curve lies above that of the active receptor.

Rewriting Eq. (51) as

$$f(x) - x = \frac{x(1-x)}{\frac{K_Y}{X_T} + x + \frac{k_3}{k_5 X_T}}.$$

shows that the deviation between the normalized dose–response curves decreases as either $K_Y/X_T$ or $k_3/(k_5 X_T)$ increases. In the limit

$$\frac{K_Y}{X_T} + \frac{k_3}{k_5 X_T} \to \infty,$$

the deviation tends to zero,

$$\lim_{\frac{K_Y}{X_T}+\frac{k_3}{k_5 X_T}\to\infty} (f(x) - x) = 0,$$

implying that the transfer function approaches the identity mapping $f(x) = x$. Therefore, perfect dose–response alignment emerges asymptotically in this limit.

However, the biological consequences of increasing these parameters differ. Increasing $K_Y$ reduces the sensitivity of the response to receptor activation, whereas increasing $k_3/k_5$ elevates basal activity, leading to constitutive activation of the downstream response protein and a progressive loss of stimulus responsiveness.

#### 1.13 Dose–response alignment when normalized by total available response protein for $k_3 > 0$

This case is similar to the previous one (i.e. when normalization was performed by $Y^*_{\mathrm{SS_{max}}}$), just that in this case, the normalized dose–response curve for the downstream response is

$$y = \frac{Y^*_{\mathrm{SS}} - Y^*_{\mathrm{SS_{min}}}}{Y_T - Y^*_{\mathrm{SS_{min}}}} = \frac{Y_T\left[\frac{X^*_{\mathrm{SS}}+\frac{k_3}{k_5}}{K_Y+X^*_{\mathrm{SS}}+\frac{k_3}{k_5}} - \frac{\frac{k_3}{k_5}}{K_Y+\frac{k_3}{k_5}}\right]}{Y_T\left[1 - \frac{\frac{k_3}{k_5}}{K_Y+\frac{k_3}{k_5}}\right]} \tag{52}$$

After simplification, this expression becomes

$$y = \frac{X^*_{\mathrm{SS}}}{K_Y + X^*_{\mathrm{SS}} + \frac{k_3}{k_5}}. \tag{53}$$

Again, $K_Y = k_4/k_5$. Defining $f(x) = y$ and substituting $X^*_{\mathrm{SS}} = X_T x$, the corresponding transfer function is

$$f(x) = \frac{X_T x}{K_Y + X_T x + \frac{k_3}{k_5}}. \tag{54}$$

The deviation from the identity mapping is therefore

$$f(x) - x = \frac{x\left[X_T(1-x) - K_Y - \frac{k_3}{k_5}\right]}{K_Y + X_T x + \frac{k_3}{k_5}}. \tag{55}$$

This expression has the same mathematical structure as that obtained for $k_3 = 0$ under normalization by total available protein. The presence of basal activity effectively replaces $K_Y$ by the combination $K_Y + k_3/k_5$. Consequently, the qualitative behavior remains unchanged.

Perfect dose–response alignment (DoRA) cannot be achieved over the entire activation range $0 \leq x \leq 1$. The two curves always intersect as $x = 0$, while a second intersection occurs at

$$x = 1 - \frac{K_Y + \frac{k_3}{k_5}}{X_T}, \tag{56}$$

provided that $K_Y + (k_3/k_5) \leq X_T$.

Rewriting the transfer function as

$$f(x) = \frac{x}{\frac{K_Y}{X_T} + x + \frac{k_3}{k_5 X_T}}, \tag{57}$$

shows that, as in the case $k_3 = 0$, the transfer function does not approach the identity mapping $f(x) = x$ in any asymptotic parameter regime. Therefore, perfect dose–response alignment does not emerge asymptotically under normalization by total available response protein.

To determine the parameter values that maximize DoRA under this normalization, we define

$$\gamma = \frac{K_Y}{X_T} + \frac{k_3}{k_5 X_T}. \tag{58}$$

The transfer function can be written as

$$f(x) = \frac{x}{\gamma + x}. \tag{59}$$

Importantly, this expression is identical in form to that obtained for $k_3 = 0$, just that in this case the term $K_Y/X_T$ is replaced by $\gamma$. Thus, the same distance measure used previously becomes

$$D(\gamma) = \int_0^1 \left| \frac{x}{\gamma + x} - x \right| dx. \tag{60}$$

Because the functional dependence on $\gamma$ is unchanged, the previous optimization holds for this general case. Numerical evaluation yields the same optimum

$$\gamma \approx 0.4622. \tag{61}$$

Substituting the definition of $\gamma$, the condition for maximal dose–response alignment becomes

$$\frac{K_Y}{X_T} + \frac{k_3}{k_5 X_T} \approx 0.4622, \tag{62}$$

or equivalently,

$$\frac{K_Y}{X_T} \approx 0.4622 - \frac{k_3}{k_5 X_T}. \tag{63}$$

Thus, unlike the case $k_3 = 0$, the optimal value of $K_Y/X_T$ is not fixed. Instead, it shifts according to the magnitude of the basal activation term $k_3/(k_5 X_T)$. Increasing basal activity therefore reduces the value of $K_Y/X_T$ required to achieve maximal dose–response alignment.

#### 1.14 Robustness of DoRA, information transmission, response range, and noise to parameter variation

##### 1.14.1 Varying receptor half-saturation constant ($K_X$)

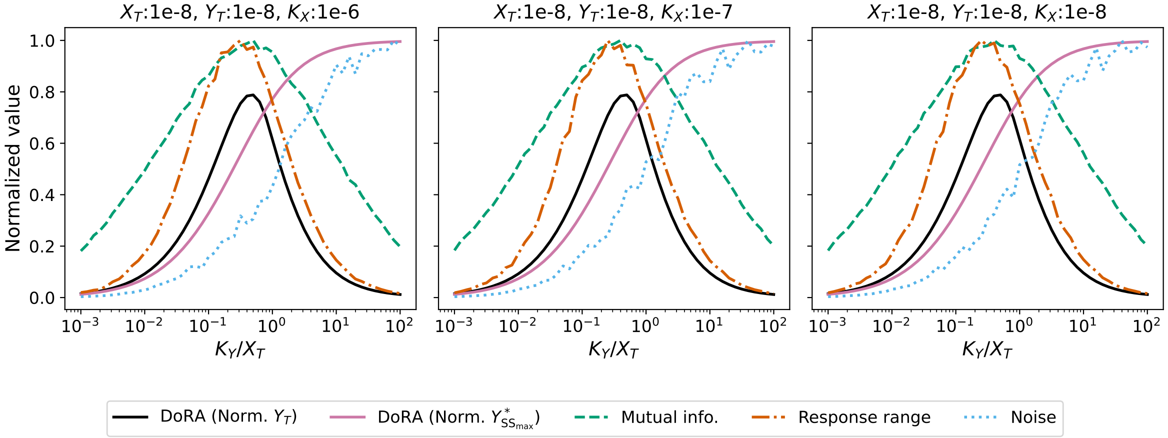


Figure S1: **Robustness of the relationship between DoRA, information transmission, response range, and noise to variations in $K_X$.** DoRA (under both normalization schemes), mutual information, response range and mean Fano factor are shown as functions of $K_Y/X_T$ for different values of $K_X$. The central panel corresponds to the parameter values used in the main text, whereas the left and right panels show the results for values one order of magnitude lower and higher for $K_X$, respectively. The overall patterns remain unchanged across the explored parameter range, indicating that the reported relationships are robust to variations in $K_X$.

##### 1.14.2 Varying total receptor ($X_T$)

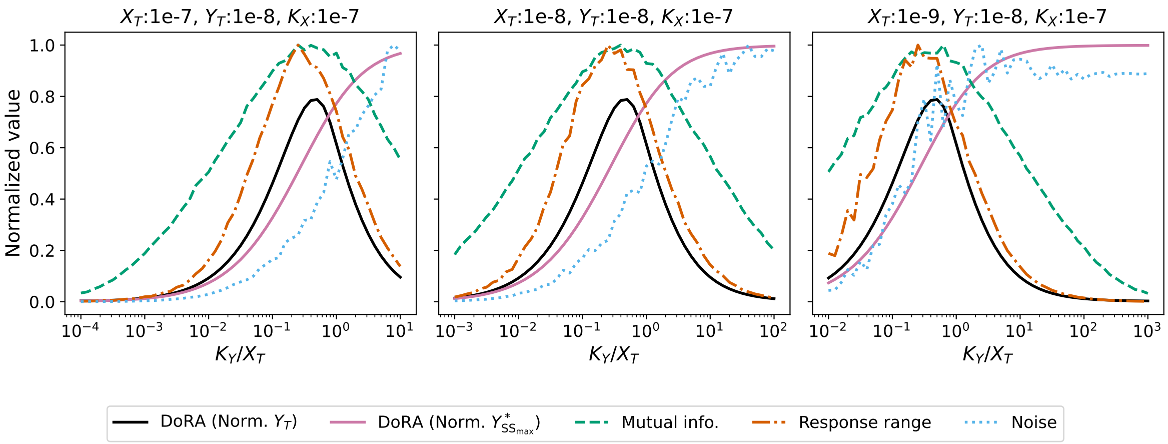


Figure S2: **Robustness of the relationship between DoRA, information transmission, response range, and noise to variations in $X_T$.** DoRA (under both normalization schemes), mutual information, response range and mean Fano factor are shown as functions of $K_Y/X_T$ for different values of $X_T$. The central panel corresponds to the parameter values used in the main text, whereas the left and right panels show the results for values one order of magnitude lower and higher for $X_T$, respectively. The overall patterns remain unchanged across the explored parameter range, indicating that the reported relationships are robust to variations in $X_T$.

##### 1.14.3 Varying total response protein ($Y_T$)

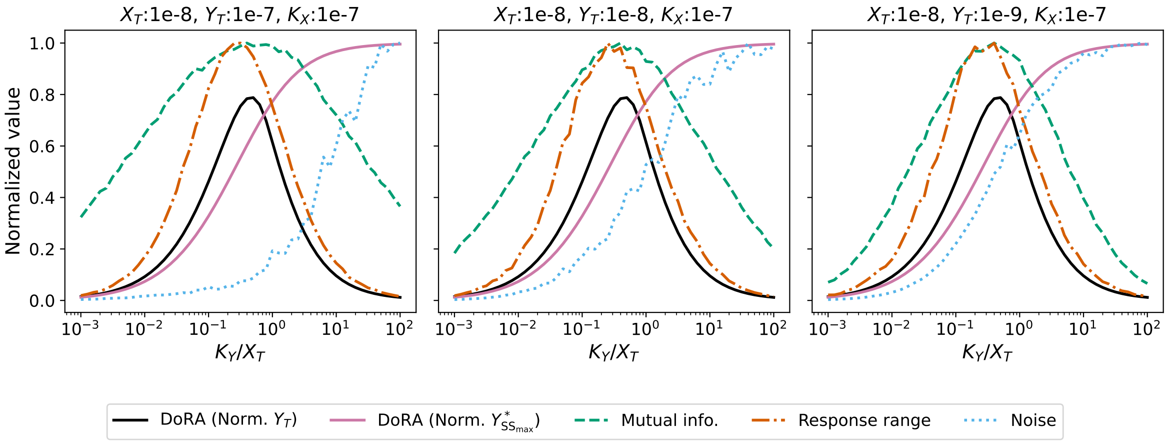


Figure S3: **Robustness of the relationship between DoRA, information transmission, response range, and noise to variations in $Y_T$.** DoRA (under both normalization schemes), mutual information, response range and mean Fano factor are shown as functions of $K_Y/X_T$ for different values of $Y_T$. The central panel corresponds to the parameter values used in the main text, whereas the left and right panels show the results for values one order of magnitude lower and higher for $Y_T$, respectively. The overall patterns remain unchanged across the explored parameter range, indicating that the reported relationships are robust to variations in $Y_T$.

### Supplementary References


[1] Khem Raj Ghusinga et al. "Molecular switch architecture determines response properties of signaling pathways". In: *PNAS* 118 (11 2021), e2013401118.

[2] Pencho Yordanov and Jörg Stelling. "Steady-State Differential Dose Response in Biological Systems". In: *Biophys. J.* 114 (3 2018), pp. 723–736.

[3] Marc Weimer et al. "The impact of data transformations on concentration-response modeling". In: *Toxicol. Lett.* 213 (2 Sept. 2012), pp. 292–298.

[4] Terry P Kenakin. *A Pharmacology Primer: Techniques for More Effective and Strategic Drug Discovery*. 6th ed. Academic Press, 2022.

[5] Lingxia Qiao, Pradipta Ghosh, and Padmini Rangamani. "Design principles of dose-response alignment in coupled GTPase switches". In: *npj Syst. Biol. Appl.* 9 (3 2023).

[6] H Kacser and J Burns. "The control of flux". In: *Symp. Soc. Exp. Biol.* 27 (1973), pp. 65–104.

[7] Reinhart Heinrich and Tom A Rapoport. "A Linear Steady-State Treatment of Enzymatic Chains: General Properties, Control and Effector Strength". In: *Eur. J. Biochem.* 42 (1 1974), pp. 89–95.

[8] Daniel T Gillespie. “A General Method for Numerically Simulating the Stochastic Time Evolution of Coupled Chemical Reactions”. In: *J. Comput. Phys.* 22 (1976), pp. 403–434.
[9] Daniel T Gillespie. “Exact stochastic simulation of coupled chemical reactions”. In: *J. Phys. Chem.* 81 (25 1977), pp. 2340–2361.
[10] Thomas M Cover and Joy A Thomas. *Elements of Information Theory.* 2nd ed. Willey-Interscience, 2006, pp. 1–748.
[11] Filipe Tostevin and Pieter Rein ten Wolde. “Mutual information in time-varying biochemical systems”. In: *Phys. Rev. E* 81 (6 2010), p. 61917.
[12] Margaritis Voliotis et al. “Information transfer by leaky, heterogeneous, protein kinase signaling systems”. In: *PNAS* 111 (3 2014), E326–E333.
[13] Ryan Suderman and Eric J Deeds. “Intrinsic limits of information transmission in biochemical signalling motifs”. In: *Interface Focus* 8 (2018), p. 20180039.
